# Chromosome-scale genome assembly of *Tinospora sagittata* (Oliv.) Gagnep. enhances identifying genes involved in the biosynthesis of jatrorrhizine

**DOI:** 10.1101/2023.07.20.549971

**Authors:** Mohammad Murtaza Alami, Shaohua Shu, Sanbo Liu, Zhen Ouyang, Yipeng Zhang, Meijia Lv, Yonghui Sang, Dalin Gong, Guozheng Yang, Shengqiu Feng, Zhinan Mei, De-Yu Xie, Xuekui Wang

## Abstract

*Tinospora sagittata* (Oliv.) Gagnep. is an important medicinal tetraploid plant in the Menispermaceae family. Its tuber, namely “Radix Tinosporae” used in Traditional Chinese Medicine, is rich in medicinal terpenoids and benzylisoquinoline alkaloids (BIAs), To enhance understanding the biosynthesis of medicinal compounds, we, herein, report the assembly of a high quality chromosome-scale genome with both PacBio HiFi and Illumina sequencing technologies. The size of assembled genome was 2.33 Gb consisting of 4070 scaffolds (N50=42.06Mb), of which 92.05% were assigned to 26 pseudochromosomes in A and B sub-genomes. A phylogenetic analysis with the *T. sagittata* and other 16 plant genomes estimated the evolutionary placement of *T. sagittata* and its divergence time in Ranunculales. Further genome evolution analysis characterized one round tandem duplication about 1.5 million years ago (MYA) and one whole-genome duplication (WGD) about 86.9 MYA. WGD contributed to the duplication of clade-specific cytochrome P450 gene family in Ranunculales. Moreover, sequencing mining obtained genome-wide genes involved in the biosynthesis of alkaloids and terpenoids. *TsA02G014550*, one candidate, was functionally characterized to catalyze the formation of (*S*)-canadine in the jatrorrhizine biosynthetic pathway. Taken together, the assembled genome of *T. sagittata* provides useful sequences to understand the biosynthesis of jatrorrhizine and other BIAs in plants.

## Introduction

*Tinospora sagittata* (Oliv.) Gagnep. is a perennial medicinal tetraploid (2n=4X=52) plant used in Traditional Chinese Medicine (TCM). It is officially listed with a Chinese medicine name “Jin Guo Lan” in the 2015 edition of the *Chinese Pharmacopoeia* (National Commission of Chinese Pharmacopoeia, 2015). To date, columbamine (Hao, 2019a), jatrorrhizine (Zhong *et al*., 2022), and palmatine (Shi *et al*., 2006) have been characterized to be three main medicinal benzylisoquinoline alkaloids (BIAs) in *T. sagittata*. In addition, Other active compounds isolated from this plant include flavonoids (D.-F. Xu *et al*., 2021), lignans (X.-Z. Huang *et al*., 2012), and clerodane type of diterpenoids (Li *et al*., 2017). These alkaloids from *T. sagittata* tuber (Radix Tinospora) have antifouling (Li *et al*., 2016), anti-inflammatory (C. Huang *et al*., 2012), and α-glucosidase inhibitory activities (Li *et al*., 2016). Moreover, these alkaloids, flavonoids, lignans, terpenoids, and other compounds provide multiple therapeutic uses of Radix Tinospora in TCM, such as improvement of immune capacity, prevention against upper respiratory infections, and lower oral ulcers, treatment of diabetes, anti-cancer, and protection of liver from different diseases (Hao, 2019b). In addition to medicinal uses, *Tinospora* is a major group of Angiosperms (Chi *et al*., 2016) and *T. sagittata* is a model plant for studying the evolutionary relationship of species within the family of Menispermaceae. It plays an important role in understanding the phylogenetic placement of Menispermaceae’s in the flowering plants.

Jatrorrhizine is a protoberberine alkaloid, a member of benzylisoquinoline alkaloids (BIAs) that are a diverse class of natural plant products (PNPs) found in numerous plants in several plant families, such as Menispermaceae, Magnoliaceae, Ranunculaceae, Papaveraceae, and Berberidaceae (Guo *et al*., 2018; Payne *et al*., 2021; Sun *et al*., 2019; Zhong *et al*., 2022). Other important medicinal BIAs found these families include berberine, palmatine, columbamine, coptisine, epiberberine, and magnoflorine (Lv *et al*., 2016). Jatrorrhizine is one of the primary bioactive metabolites of *T. sagittata* (Zhong *et al*., 2022). It has antidiabetic (Zhong *et al*., 2022), antimicrobial (Ali *et al*., 2013), antiprotozoal (Malebo *et al*., 2013), anticancer (Sun *et al*., 2019), anti-obesity and hypolipidemic (Yang *et al*., 2016) activities. In addition, this compound benefits central nervous system activities (Bacq *et al*., 2012). To date, although the full biosynthetic pathway of jatrorrhizine remains open for studies, previous studies have suggested that it derives from L-tyrosine through the ring-opening of berberine (Beecher and Kelleher, 1983; Rueffer *et al*., 1983). Particularly, the berberine biosynthesis investigated in *Coptis chinensis* (Liu *et al*., 2021) and *Papaver somniferum* (Guo *et al*., 2018) has provided important gene and enzyme information that can enhance understanding the formation of jatrorrhizne in *T. sagittata*. Meanwhile, an integration of genomics, transcriptomics, and metabolomics is helpful to expedite understanding the biosynthesis of jatrorrhizine. Herein, to enhance understanding the genome features of *T. sagittata* and the biosynthesis of jatrorrhizine and other medicinal compounds in this medicinal plant, we report genome and transcroptome assembly with different sequencing technologies. Meanwhile, we completed both targeted and non-targeted metabolic profiling for characterization of BIAs and other plant secondary metabolites in this medicinal plant. We assembled a high quality genome and several transcriptomes that allowed characterizing the phylogenetic placement of *T. sagittata* in the Menispermaceae family and of the divergence time of this family in Ranunculales. The genome of *T. sagittata* provides fundamental knowledge to study the evolutionary and integrative biology of Ranunculales. Furthermore, the integration of genomes, transcriptomes, and metabolic profiling disclosed candidate genes associated with the biosynthesis of jatrorrhizine and other BIAs in this medicinal plant.

## Method and materials

### Plant Materials

*Tinospora sagittata*, a medicinal tetraploid plant, was cultivated for its rhizomes, known as Radix Tinosporae (RT). Seeds from Lichuan, Hubei Province, China, were planted in controlled growth conditions. After four months, seedlings were transferred to Huazhong Agricultural University for further research, and young and healthy leaves were collected for DNA extraction, gene expression profiling, and metabolomics.

### Genome sequencing with PacBio technology

Genomic DNA was extracted from fresh leaves using the DNAsecure Plant Kit, followed by shearing the DNA to 10 kb size using the Megaruptor® DNA Shearing System. SMRTbell libraries with 60 kb insert size were constructed from at least 10 μg of sheared DNA using the SMRTbell Express Template Prep Kit 2.0. Hi-C libraries were prepared by cell cross-linking, endonuclease digestion, end repair, circularization, and DNA purification and capture (Xie *et al*., 2015). The libraries’ concentration and insert size were determined, and the Illumina platform was used for paired-end sequencing with a read length of 150 bp.

### Genome assembly and quality assessment

The *T. sagittata* genome size was estimated using a *k*-mer analysis approach and Illumina PE short reads. Long reads from the PacBio SMRT Sequencer were assembled using FALCON (Chin *et al*., 2016). Error correction, contig assembly, and polishing were performed. Errors in the Illumina sequencing reads were corrected using Pilon (Walker *et al*., 2014). Hi-C sequencing results were aligned to the assembled scaffolds. The scaffolds were then clustered into chromosomes using LACHESIS (Burton *et al*., 2013). The quality of the drafted genome was evaluated using various tools, including read mapping, transcriptome mapping, CEGMA (Parra *et al*., 2007), and BUSCO (Simão *et al*., 2015) analysis. These assessments aimed to determine genome assembly, gene set, and transcriptome completeness.

### Genes annotation

The gene function annotation process involved three main methods: homology-based prediction, de novo prediction, and transcriptome-based prediction. Homologous proteins from four plant genomes were obtained and aligned to the *T. sagittata* genome using TblastN (Altschul *et al*., 1990). The resulting BLAST hits were combined using Solar software (Yu *et al*., 2006), and GeneWise (Birney *et al*., 2004) was used to predict the gene structure. For transcriptome-based prediction, RNA-seq data was mapped to the genome and assembled into gene models using TopHat (Trapnell *et al*., 2009) and Cufflinks (Trapnell *et al*., 2010). Additionally, RNA-seq data was assembled with Trinity (Grabherr *et al*., 2011) to create pseudo-ESTs, which were then mapped to the genome for gene prediction using PASA.

Ab initio gene prediction programs were employed to predict coding regions in the repeat-masked genome. The gene structures obtained from different prediction methods were combined using EvidenceModeler (EVM) (Haas *et al*., 2008) to generate a non-redundant set of gene structures. Protein sequences were subjected to BLASTP (Gish and States, 1993) against SwissProt and NR databases, and protein domains were annotated using InterPro (Hunter *et al*., 2009) and Pfam (Mistry *et al*., 2021) databases. Function prediction of protein-coding genes utilized InterProScan (Quevillon *et al*., 2005) and HMMER (Finn *et al*., 2011).

Gene Ontology (GO) terms were obtained from InterPro or Pfam entries. Genes associated with jatrorrhizine biosynthesis were identified by BLAST against KEGG databases. tRNA genes were identified using tRNAscan-SE (Lowe and Eddy, 1997), while rRNA fragments were predicted by aligning transcripts to rRNA sequences. cDNAs encoding miRNA and snRNA were predicted using INFERNAL (Nawrocki *et al*., 2009) against the Rfam database.

The identification of transposable elements (TEs) involved a combination of de novo and homology-based approaches. RepeatModeler, LTR_FINDER, and RepeatScout were used to build a repeat library through the de novo approach. The homology-based approach utilized RepeatMasker against the Repbase TE library and RepeatProteinMask against the TE protein database. Tandem repeats were detected using the Tandem Repeats Finder (TRF) software (Benson, 1999).

### Gene family analysis

The online tool plantiSMASH (Kautsar *et al*., 2017) was used to analyze biosynthesis cluster genes (BCGs). Homologs were searched in the *T. sagittata* genome using BIA biosynthesis genes from *C. chinensis* as queries. *A. thaliana* and rice CYP gene sequences were used to search for homologs. Genes <300 amino acids were removed. ClustalW and MEGAX were used for multiple sequence alignment and tree construction (Kumar *et al*., 2018).

### Phylogenomic analysis

Gene families were clustered, and MUSCLE (Edgar, 2004) was used for aligning single-copy gene protein sequences. A super alignment matrix was created from the combined results. A phylogenetic tree was constructed using RAxML(Stamatakis, 2014) with *T. sagittata* and 16 other species, and MCMCtree (Yang, 2007) in PAML was used to infer divergence time based on the tree.

### Identifying expansion and contraction of gene families

We used the CAFÉ program (De Bie *et al*., 2006) to analyze the expansion and contraction of gene families through comparing the cluster size differences between the species. A random birth and death model was used to study changes in gene families along each lineage of the phylogenetic tree. A probabilistic graphical model (PGM) was introduced to calculate the probability of transitions in the gene family size from parent to child nodes in the phylogeny. We calculated the corresponding p-values in each lineage using conditional likelihoods as the test statistics. A *p*-value of <0.05 was used to identify significantly expanded and contracted families.

### Analysis of whole-genome duplication

The study utilized the 4DTv and *Ks* methods to analyze whole genome duplication (WGD) events in *T. sagittate* (Liu *et al*., 2021; Zwaenepoel and Van de Peer, 2019). Syntenic blocks were constructed by comparing protein sequences from *T. sagittata*, *C. chinensis*, *Papaver somniferum*, *Vitis vinifera*, and *Tripterygium wilfordii* using BLASTp(E<1e^-5^). MCScanX (Tang *et al*., 2008) identified syntenic blocks with at least five genes. 4DTv (Tu *et al*., 2020) and *Ks* values were calculated for syntenic segments to understand their distribution. The wgd v1.1.1 software (Zwaenepoel and Van de Peer, 2019) determined *Ks* values, which were converted to divergence time using a substitution rate. The mean *Ks* value and standard deviations of WGD duplications were calculated.

### Analysis of genome synteny

The study utilized MUMmer 4.0 (Marçais *et al*., 2018) for genome alignment and SyRI for variant detection, including collinearity regions and structural rearrangements. Identified mutations in gene regions were annotated with GO, KEGG, and other databases using BLAST. GO and KEGG enrichment analyses were conducted with clusterProfiler v3.14.0 (Yu *et al*., 2012).

### RNA isolation from different *T. sagittata* tissues and Illumina sequencing

Plant tissues (leaves, stems, roots, and tubers) were harvested from greenhouse-grown plants, frozen in liquid nitrogen, and stored at -80 °C. Total RNA was extracted using a modified CTAB method (Allen *et al*., 2006). RNA samples were treated with DNaseI, evaluated on agarose gel and Bioanalyzer, and used for cDNA library construction. Three libraries per tissue were sequenced on an Illumina HiseqX platform, generating 150-bp paired-end reads (Mendelevich *et al*., 2021).

### Assembly of transcriptomes and transcriptional analysis

RNA-seq reads were filtered and trimmed to obtain high-quality clean reads. TopHat2 (Kim *et al*., 2013) was used to map the reads to the genome, and HTSeq (Anders *et al*., 2015) provided Reads Per Kilobase per Million (RPKM) values for each gene. DESeq2 (Love *et al*., 2014) normalized gene expression, and differentially expressed genes (DEGs) were identified based on adjusted *p*-values. GOseq (Young *et al*., 2010) analyzed DEG enrichment and annotation, while KEGG categorized genes and KOBAS (Xie *et al*., 2011) assessed enrichment in KEGG pathways. Candidate genes involved in jatrorrhizine biosynthesis were identified from transcriptomes, and their expression profiles were analyzed in different plant parts for correlation with jatrorrhizine levels measured by HPLC.

### Measurement of jatrorrhizine and metabolic profiling

According to our previous report (Deng *et al*., 2008), high-performance liquid chromatography (HPLC) was performed to measure jatrorrhizine in different tissues. Meanwhile, ultra-performance liquid chromatography quadrupole time of flight mass spectrometry (UPLC/Q-TOF MS) was used to profile metabolites according to our previous report (Tang *et al*., 2008) Both targeted and untargeted metabolite levels were used as matrix for PCA analysis to understand the differentiation among different tissues.

### Quantitative real-time PCR analysis

Total RNA was extracted using the FastPure Universal Plant Total RNA Isolation kit (Vazyme) by following the manufacture’s protocol and then was treated with DNAase to remove genomic DNA. Two μg of DNA-free RNA was used as the template for reverse transcription to synthesize the first-strand cDNA using the HiScript II Q RT SuperMixfor qPCR kit (Vazyme) by following the manufacture’s instructions. Two µl of cDNA was used as the template for quantitative real-time PCR in 20-µl volume with 2x M5 HiPer SYBR premix Es Taq (Mei5 Biotechnology, Co., L, China). A pair of primers (Table S23) were designed to amplify *TsA02G014550*. A pair of primers were designed to amplify *ACTIN* a housekeeping gene as reference (Table S24). The reaction condition was composed of 95°C for 30s for 1 cycle followed by 95°C for 5s, 60°C for 45s for 40 cycles, and then 72°C for 20s. The relative expression level of *TsA02G014550* was calculated using 2^-ΔΔCt^ method.

### Heterologous expression of *TsA02G014550* in yeast and in vitro activity assay

The open reading frame (ORF) of *TsA02G014550* was cloned to an epitope-tagged pESC-His vector for heterologous expression according to a reported protocol (Liu *et al*., 2021). An epitope-tagged pESC-His vector fused with *TsA02G014550* gene, namely as pESC-TsA02G014550, was constructed to express its recombinant protein. The pESC-TsA02G014550 vector was transformed into the yeast strain WAT11 that expressed a modified endogenous NADPH-CYP reductase. In addition, the pESC-His vector was introduced into the same strain of yeast as a control. Yeast cultures were placed in 50 ml tubes and centrifuged for 10 min at 5000 rpm. The pellets were resuspended twice in respective TEK and TESB buffers and were then broken up with a cryogenic homogenizer. Further centrifugation was conducted for the precipitation of microsome pellets. For in vitro activity assays, 0.5 mg resuspended microsomal protein and 100 μM of a potential substrate were included in a 500 μL reaction system to incubate possible products. Alkaloids from catalytic products were identified using an ultra-high performance liquid chromatography-quadrupole time-of-flight mass spectrometry system (Waters, Milford, MA, USA) with elution buffers consisting of 0.1% aqueous formic acid and 0.1% formic acid in acetonitrile. MS analysis was conducted on a Waters XevoG2-S QTOFmass spectrometer with electrospray ionization. The acquisition of data was controlled by Waters MassLynx software (v4.1).

## Results

### Genome sequencing and assembly

A 21-*mer* analysis of the sequenced genome revealed that allotetraploid *T. sagittata* had a monoploid genome size of ∼553.23 Mb and a whole-genome size of 2.33 Gb. This analysis indicated that the genome of *T. sagittata* was characterized with a 2.98% heterozygosity. Although the heterozygosity challenged our de novo assembly, we estimated that there were 4,328,940 biallelic heterozygous sites throughout the 26 assembled chromosomes. We also characterized the average heterozygosity that corresponded to one heterozygous single-nucleotide polymorphism (SNP) per 212 bp (Fig. 1B and Table S1).

**Fig. 1.**
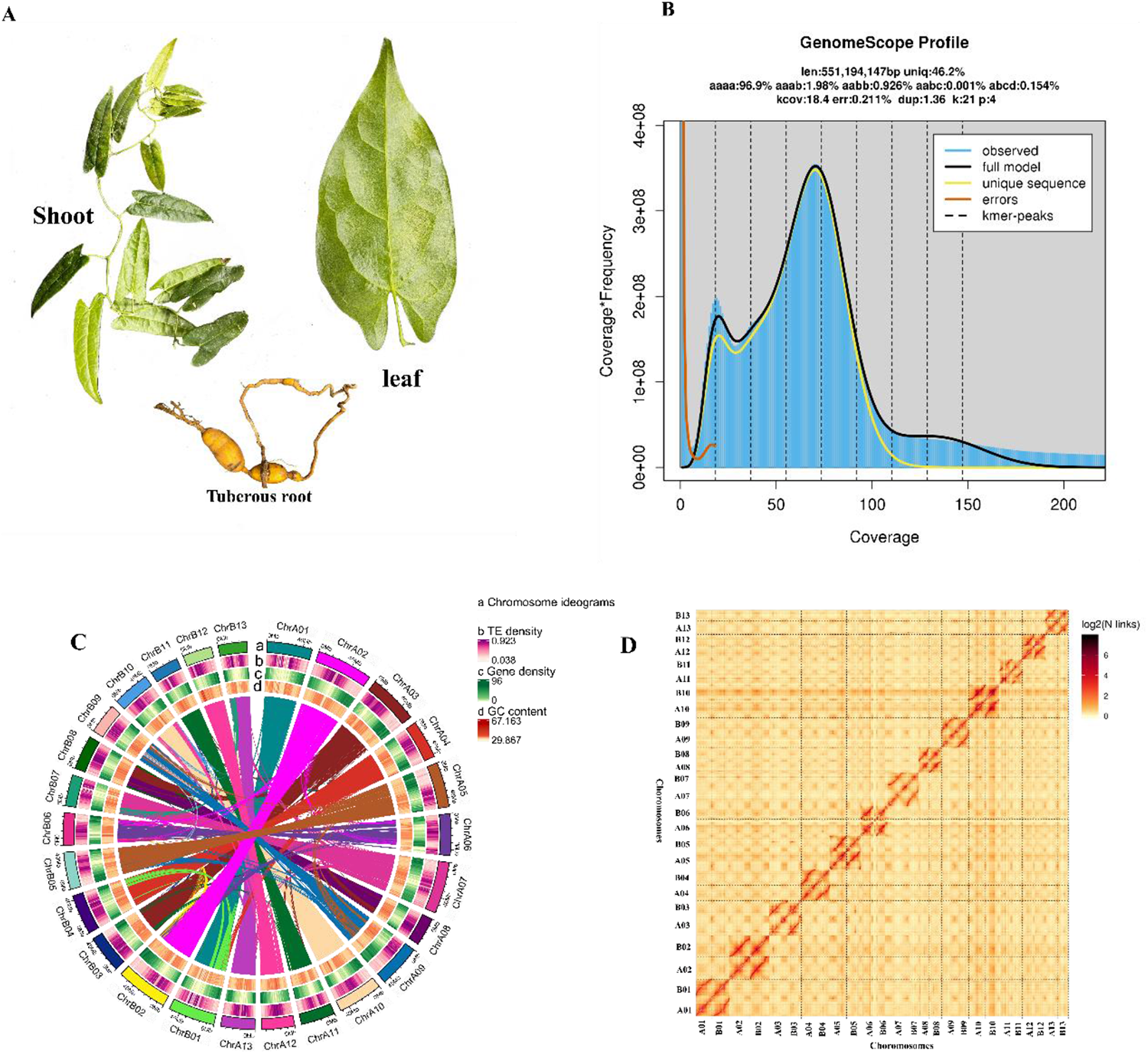
Main features of genome assembly of *T. sagittata*. (A) Three types of tissues of *T. sagittata* were collected from a research station at Huazhong Agricultural University, Hubei Province, China. (B) *21-mer* depth distribution of sequencing reads. (C) Distribution features of the assembled genome are shown by four types of elements arranged from outer (a) to inner (d), (**a**) chromosomes ideograms, (b) transposon elements (TE) density, (**c**) gene density, and (d) GC content. (D) An interaction heat map of Hi-C assembled chromosomes.

The sequencing with the PacBio Sequel II yielded 2.5 million CCS reads (average length 15.7kb) with a total data volume of 43 GB. The PacBio long reads were corrected and assembled with a hybrid assembly strategy prior to using 11.25 Gb of Illumina sequencing (short reads) for polishing (Fig. S1&2, Table S2-5). The assembly of short reads from the Illumina sequencing and PacBio long reads resulted in 4483 contigs (N50=8.3Mbp) with a total of 1299.6 Mbp, in which the maximum length was 31.9Mbp and the GC content was 36.36% (Table 1).

**Table 1.** Statistical summary of the genome assembly of *T. sagittata*.

| **Data source** | **PacBio+Illumina** | **PacBio +Hi-C** |
| --- | --- | --- |
| Total scaffolds |  | 1,299,652,234 |
| Number of scaffolds |  | 4,070 |
| Scaffold N50(bp) |  | 42,067,207 |
| Scaffold N90(bp) |  | 66,947 |
| Scaffold Max |  | 63,028,082 |
| Total contigs Length | 1,142,509,881 | 1,299,610,934 |
| Number of contigs | 2,531 | 4,483 |
| Contig N50(bp) | 23,237,671 | 8,349,146 |
| Contig N90(bp) | 1,234,648 | 65,341 |
| Contig Max | 49,008,001 | 31,913,866 |
| GC % | 36.49 | 36.36 |

After the polishing, PacBio long reads with Illumina short reads the assembly was further scaffolded with Hi-C. The scaffolding results obtained 4070 scaffolds (N50=42.06Mb). Subsequently, 960.8 million reads with 287,909.2 Mbp clean Hi-C paired-end reads were used for scaffold extension and anchoring. The Hi-C assembly and manual adjustment of the heatmap obtained 1,196.2 Mbp of genomic sequences accounting for 92.05%, which were used for mapping to the 26 chromosomes. The results showed that 1,132.3 Mbp out of 1,196.2 Mbp (accounting for 94.66%) were mapped to the 26 chromosome sequences. A further sequence analysis obtained 572.7 Mb of reads that were uniquely aligned to the genome. In these unique sequences, 399.5 Mb (accounting for 69.75% of the uniquely aligned reads) were valid Hi-C data visualized with a heat map (Fig. 1D, Table 1, Table S6 and S7). In the heat map (Fig. 1 D), the 26 chromosomes were clearly distinguished to form 4070 unique groups. In each group, the intensity of the interaction at its diagonal position was higher than that at the non-diagonal position, indicating that those chromosomes assembled by Hi-C were adjacent to each other. The heat map also showed that the interaction signal strength between the sequences at the diagonal position) were strong, while that between non-adjacent sequences at off-diagonal position was weak (Fig. 1D, Fig. S3). This result is consistent with the principle of Hi-C-assisted genome assembly. Based on the assembly of the chromosome-based genome, the analysis with the Circlize software provided ideograms, gene density, repeat density, GC content, and collinearity between the chromosomes (Fig. 1C a-d).

The Illumina short reads were aligned to the assembled genome by using BWA (Li and Durbin, 2009) to evaluate the quality of the assembly. The results revealed that the mapping rate of illumine and PacBio sequencing was about 99.27%. Then, based on plant gene models, Benchmarking Universal Single-Copy Orthologs (BUSCO) analysis was completed to quantitatively assess the assembled genome. The results indicated that 96.78% of the BUSCO sequences were present in the *T. sagittata* genome, while only 0.99% and 2.23% of them were fragmented and missing, respectively (Table S8). Furthermore, Core Eukaryotic Genes Mapping Approach (CEGMA) analysis (Parra *et al*., 2007) was completed to understand core protein-encoding orthologs. The resulting data disclosed that in 458 core eukaryotic genes (CEG), 421 (about 91.92% of CEGMA) were presenting in the assembled *T. sagittata* genome. In addition, in 248 highly conserved CEGs, 190 (about 76.61%) existed in the assembled genome (Table S9). Based on similar analysis in recently reported *opium poppy* (Guo *et al*., 2018) and *T. wilfordii* (Tu *et al*., 2020) genomes, these data indicated a highly accuracy and nearly completeness of the *T. sagittata* genome assembled here. Taken together, this assembly provided a *high*-quality chromosome-level genome of *T. sagittata*.

### Genome annotation

Three methods, homology-based, Ab-initio-based, and RNAseq-based, were used to predict protein-coding genes. After the removal of theoretical nonfunctional genes, 52,953 protein-coding genes was obtained from the assembled genome (Table S10). Among the predicted genes, 1047, 3788 and 10 genes were unique in homology-based, Ab-initio-based, and RNAseq-based, respectively (Fig. S4).

Meanwhile, tissue-specific RNA-seq was completed to develop transcriptomes. The resulting data showed that the average length of coding sequence genes was 6203.59 bp. The length of average coding sequence (CDS) were 1360.42 bp, with an average of five exons and four introns per gene (Table S11). Approximately 97.93% of the genes were functionally annotated, of which 96.91 % and 97.79% had significant hits in the NR and TrEMBL databases, respectively. Gene Ontology (GO) terms classified 82.91 % of the genes. The use of Kyoto Encyclopedia of Genes and Genomes (KEGG) pathways annotated 75.21 % of the genes (Table S12). These results indicate the high accuracy of the gene predictions in the *T. sagittata* genome. We further annotated noncoding RNA, yielding 9,624 transfer RNA (tRNA) genes, 13,014 ribosomal RNA (rRNA) genes, 350 small nuclear RNA genes, and 292 microRNA genes, as well as 287 pseudogenes in the *T. sagittata* genome (Table S13-15).

A total of 1,119,004 (51.72%) reads with a length of 672.2 Mb of the assembly was masked and annotated as repetitive elements, of which 45.13% was retroelement, while 6.59% was DNA transposon. Of the repetitive elements, long terminal repeat (LTR) counted for 41.85%, and long interspersed nuclear elements (LINE) were 2.92%. Interestingly, most LTRs were *Gypsy* and *Copy* elements (constituting 18.55% and 14.39% of the *T. sagittata* genome), and 8.32% comprised unknown LTR repeats (Table S16). About 91 Mb of tandem repeats were obtained, accounting for 7% of the genome (Table S17).

We next combined RNA-seq and full-length transcriptome data from four different tissues and organs (leaf, rhizome, roots, and stem) with three biological replicates. At least 6.36 Gb of clean data were generated for each sample, with a minimum of 94.02% clean data achieving a quality score of Q30 (Data S1). Clean reads of each sample were mapped to a specified reference genome. The mapping ratio ranged from 89.11% to 93.96% (Table S18). Prediction of alternative splicing, gene structure optimization analysis, and novel gene discovery was processed on top of mapping results, during which 4,905 were discovered, and 2,215 novel genes were annotated with a putative function.

Increased amounts of evidence have shown that plant alkaloid accumulation is regulated by transcription factors (TFs) from the WRKY, bHLH, or AP2/ERF superfamilies. With the assembled *T. sagittata* genome, we identified 134 WRKY, 214 bHLH, and 230 AP2/ERF genes in total, and these genes fell into diverse subfamilies, respectively (Data S10).

### Genome comparison of *T. sagittata* and 16 other species

To understand the evolution of the *T. sagittata* genome, a comparative analysis was performed using 16 other plant genomes (Table S19). The sequence homology blast in these 17 genomes obtained 625,332 protein-coding genes (including 515,061 genes in ortho groups and 110,271 unassigned genes), 58,911 of which were obtained in the *T. sagittata* genome (Fig. S5, Data S2). Gene family analysis revealed that 58,911 genes consisted of 2248 families. Further sequence mining of the 625,332 genes showed that there were 1518 gene families shared by the 17 species. Accordingly, this analysis indicated that *T. sagittata* had 766 unique gene families. In addition, of the other 16 plant species, the genome of *C. chinensis* was found to be the closest to that of *T. sagittata*. Two have 181 gene families in common (Fig. 2A and B). A synteny analysis was completed to compare the genomes of *T. sagittata*, *C. chinensis*, *P. somniferum*, *T. wilfordii*, and *V. vinifera*. The results showed 30,085 homologous genes (about 32.36% of the total number of genes, TNGs) between *T. sagittata* and *C. chinensis*, 42,397 homologous genes (about 36.59% of TNGs) between *T. sagittata* and *P. somniferum*, 39, 038 homologous genes (about 46.26% of TNGs) in *T. sagittata* and *T. wilfordii*, and 34,435 homologous genes (43.42% of TNGs) between *T. sagittata* and *V. vinifera* (Fig. 2C, Fig. S6, Table S20).

**Fig. 2.**
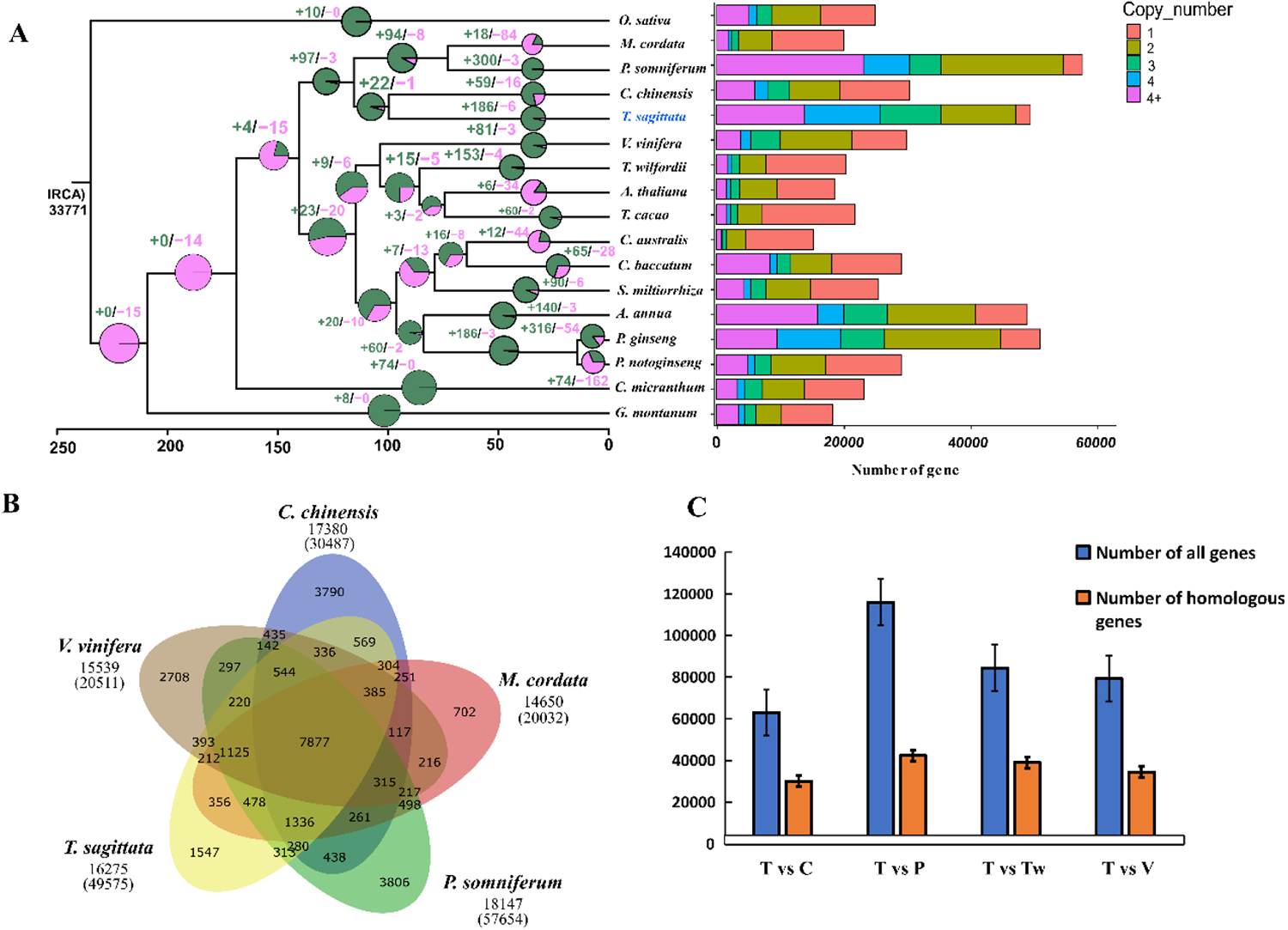
Comparative analysis of *T. sagittata* and sixteen other plant genomes. (A) A phylogenetic tree and a histogram were developed from *T. sagittata* and 16 other flowering plants. Expansion and contraction of gene families are denoted with numbers highlighted by a plus (+) and a minus (-) sign, respectively. All the nodes are 100% bootstrap support (left panel), and the histogram shows the number of genes distributed in each species resulted from the gene family classification (right panel). (B) A Venn diagram showed gene family clustering. The total numbers of gene families and the total number of genes in parentheses under each species name are provided. The number in the Venn diagram means gene families. (C) The collinearity ratio between *T. sagittata* and each of four other species were estimated. T; *T. sagittata,* C; *C. chinensis*, P; *P. somniferum,* Tw; *T. wilfordii,* and V; *V. vinifera*.

### Functional categorization of unique gene families

GO and KEGG enrichment analyses were performed to categorize molecular functions of the 766 unique gene families. The resulting data of GO enrichment indicated that those gene families were associated with RNA-DNA hybrid ribonuclease activity (GO: 0004523), regulatory region nucleic acid binding (GO: 0001067), transcription regulatory region sequence-specific DNA binding (GO: 0000987), and cis-regulatory region sequence-specific DNA binding (GO: 1900750) (Fig. S7, Data S3). Meanwhile, the KEGG analysis showed that the unique gene families were enriched in benzoxazinoid biosynthesis (ko00402), ABC transporters (ko02010), monoterpenoid biosynthesis (ko00902), and mRNA surveillance pathway (ko03015) (Fig. S8, Data S4).

### Expansion of gene families, phylogenetic features, and whole genome duplication

The 2284 gene families were used for gene family evolution analysis. The results indicated that 186 were expanded and 6 were contracted in the *T. sagittata* genome (Fig. 2A). The GO enrichment analysis showed that the 186 expanded families were related to DNA helicase activity (GO: 0003678), UDP-glycosyltransferase activity (GO: 0008194), ADP binding (GO: 0043531), and RNA-DNA hybrid ribonuclease activity (GO: 0004523) (Fig. S9, Data S5). The KEGG analysis indicated that the 186 expanded gene families involved in benzoxazinoid biosynthesis (ko00402), 2-oxocarboxylic acid metabolism (ko01210), sesquiterpenoid and triterpenoid biosynthesis (ko00909), and isoquinoline alkaloid biosynthesis (ko00950). This enrichment analysis revealed that Bias bridge enzyme-like 18, mdis1-interacting receptor-like kinase 2-like, cytochrome p45071D10, BNAA01g32320d protein, OS12G0491200 protein, and CME8 protein gene families were expanded in *T. sagittata* genome (Fig. S10, Data S6). More importantly, the expanded gene families included CYP and BBE genes that were likely associated with the biosynthesis of jatrorrhizine.

To investigate the phylogenetic placement of *T. sagittata*, phylogenetic trees were constructed from 17 species. We completed an ortholog analysis, obtained 1113 (defined: 41.2% of species having single-copy) single-copy genes from the *T. sagittata* and 16 other medicinal plant genomes (Fig. S5), and then used them for phylogenetic analysis. The single-copy genes were used to construct a species tree through maximum likelihood analysis of a concatenated supermatrix. The results from the two analyses indicated that their consistent topographies clustered *T. sagittata* as a sister lineage of *C. chinensis*. It was interesting that these two species were closely related to *Macleaya cordata* and *Papaver somniferum* L. In addition, the divergence times of these plants were estimated using MCMCtree with calibrations (Fig. 3D). The resulting data indicated that the divergence time of *T. sagittata* and *C. chinensis* was approximately 89-109 MYA during the Cretaceous period. In addition, the data estimated that two diverged from the *Macleaya cordata* and *P. somniferum* L. approximately 108-120 MYA.

**Fig. 3.**
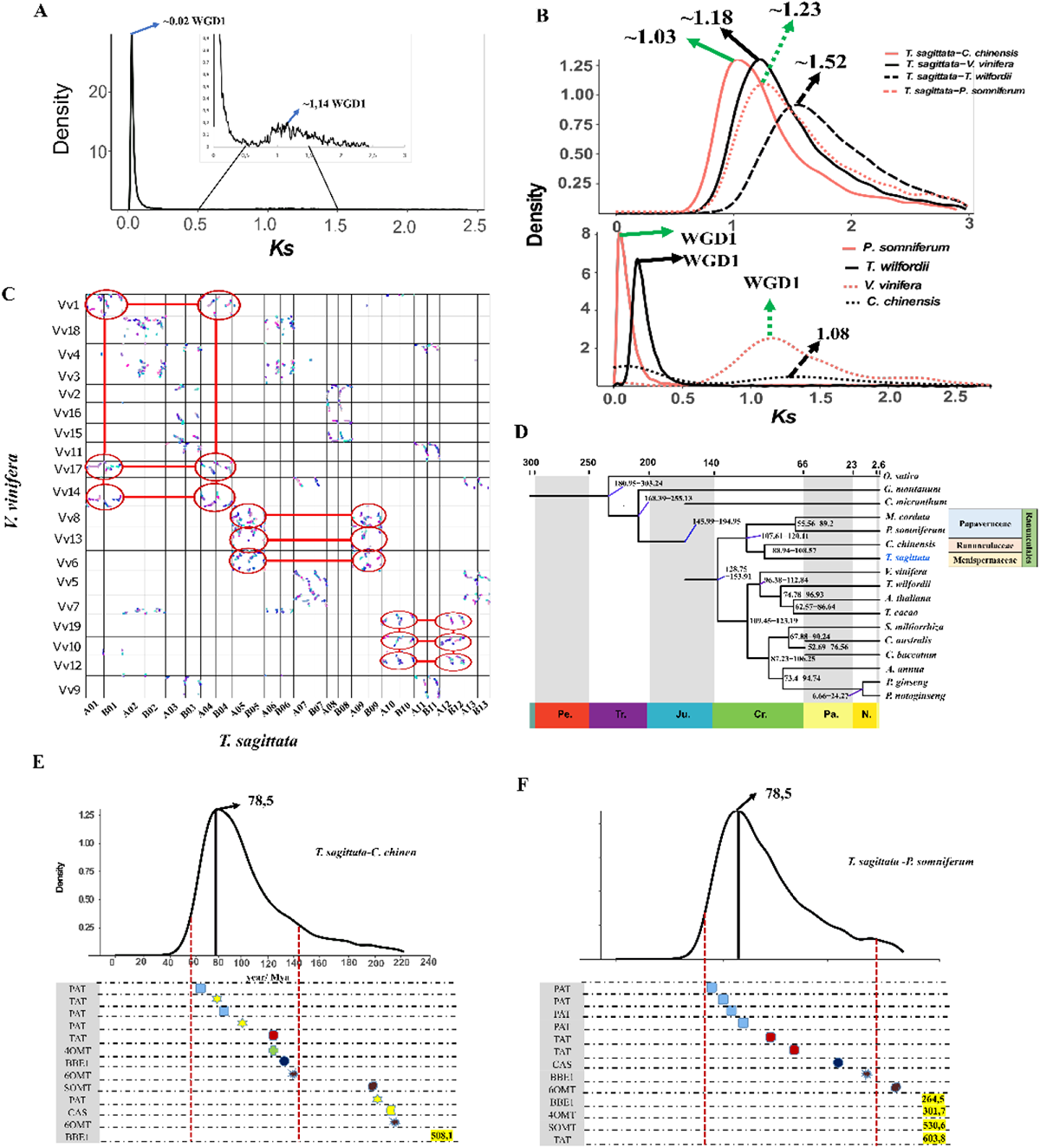
Evolutionary analysis of *T. sagittata* and other 16 angiosperm genomes as well as mining of genes involved in the jatrorrhizine biosynthesis. (A) synonymous divergence (*Ks*) for orthologue duplicates identified within *T. sagittata* (both transcriptome and genome data were used). (B) Density plots show synonymous substitution rate (*Ks*) distributions of syntenic blocks for *T. sagittata* with *C. chinensis, V. vinifera, T. wilfordii,* and *P. somniferum* (top). Density plots show the synonymous substitution rate (*Ks*) distributions in *C. chinensis, V. vinifera, T. wilfordii,* and *P. somniferum* (bottom). (C) Syntenic dot plots show a 3:1 chromosomal relationship between the *T. sagittata* genome and *V. vinifera* genome. (D) An evolutionary tree was developed with differentiation times. The numbers indicate the estimated divergence time of each node in millions of years. (E) The duplication time was estimated for 3 jatrorrhizine biosynthesis genes in the genome of C*. chinensis*, (F) The duplication time was estimated for 13 jatrorrhizine biosynthesis genes in the genome of *P. somniferum*.

Paralogous genes in the genomes of *T. sagittata*, *V. vinifera, T. wilfordii, P. somniferum,* and *C. chinensis* were identified to characterize whole genome duplication (WDG). The WGD events were examined via distributions of synonymous substitutions (*Ks*) within genes in syntenic blocks. The results showed that the distribution of *Ks* for the paralogous genes of the *T. sagittata* genome had two prominent peaks at ∼0.02 (1.5 MYA) and ∼1.14 (86.9 MYA). These two peak times indicated that the genome of *T. sagittata* experienced one round tandem duplication and the other round of ancient WGD events (Fig. 3A). The same analysis was done with the genomes of two other species of Ranunculales, *P. somniferum* (Guo *et al*., 2018) and *C. chinensis* (Liu *et al*., 2021), revealed the occurrence of WGDs event in *T. sagittata* and *P. somniferum* (Fig. 3B). In addition, the distribution of fourfold degenerate synonymous sites of the third codons (4DTv) of all gene pairs found in each segment showed two peaks at approximately 0.01 and 0.33 in the *T. sagittata* genome (Fig. S11). The relative age (Kimura distance) computed for LTR retroelements also indicated a recent increase of transposon activity (Fig. S12). The most recent outbreak of LTRs in the *T. sagittata* genome might be associated with the the large genome size.

Furthermore, we comparatively investigated the WGD events of *T. sagittata* vs. *C. chinensis* and *T. sagittata* vs. *P. somniferum*, while those of *T. sagittata* vs. *T. wilfordii* and *T. sagittata* vs. *V. vinifera* were used as the outgroups. The peak distribution of *Ks* values for *C. chinensis*, *P. somniferum, T. wilfordii,* and *V. vinifera* were *Ks* ∼1.03, 1.23, 1.52, and 1.18, respectively (Fig. 3B). In addition, an analysis of the syntenic dot plots was completed to understand ancient WGDs in the *T. sagittata* genome. The resulting data indicated that a 2:3 syntenic depth ratio between *T. sagittata* and grapevine (*V. vinifera*) was triplicated during the eudicot hexaploidy event (Jaillon *et al*., 2007) (Fig. 3C). Furthermore, Based on the reports that the *C. chinensis* and *P. somniferum* genomes experienced one WGD after the ancestral angiosperm genome duplication event (Guo *et al*., 2018; Liu *et al*., 2021), the synteny relationship between *T. sagittata* and *C. chinensis* and *P. somniferum* (Fig. S13-14) was compared to understand ancient WGD events in *T. sagittata.* This comparison obtained a 2:2 syntenic depth ratio, suggesting that the genome of *T. sagittata* experienced only a single ancient WGD.

In addition, we calculated the *Ks* values for each duplicated gene pair associated with BIA biosynthesis in *T. sagittata* vs. *C. chinensis* and in *T. sagittata* vs. *P. somniferum*. The resulting data revealed major duplications of related genes in the biosynthetic pathway occurred in the recent WGD events (Fig. 3E and F). These data suggested that the recent WGD events were associated with the evolution of the BIA biosynthesis in *T. sagittata* (Table S21-22).

### Structure differences between A and B sub-genomes

We conducted a collinearity analysis of chromosomes to determine the syntenic relationship between A and B sub-genomes. The results showed that each of the 13 chromosomes in the A sub-genome had their apparent collinear chromosome in the B sub-genome (Fig. 4), indicating that the sub-genomes originated from the same ancestor. Based on these features, the A sub-genome was selected as the reference for analyzing the syntenic regions, translocation, inversion, and duplication variation in the B sub-genome. This analysis determined 2311 chromosomal translocations, 208 inversions, and 6334 duplications in the B sub-genome (Fig. S15). Furthermore, analyzed were the single-nucleotide polymorphisms (SNPs), small insertions and deletions (InDels), and structure variation (SVs) in aligned syntenic blocks in the B sub-genome (Fig. S16). This analysis unearthed 3.5 million SNPs, 877,086 InDels (556,367 insertions, 320,719 deletions), and 8,853 SVs that were identified between the A and B sub-genomes. Finally, an analysis of the presence and absence variation (PAV) of distributions across the chromosomes obtained 9,166 PAVs with a length of 9.1 Mb in the B sub-genome (Fig. S17). Moreover, the PAV analysis detected 688 and 683 copy gains and losses in the B sub-genome, respectively.

**Fig. 4.**
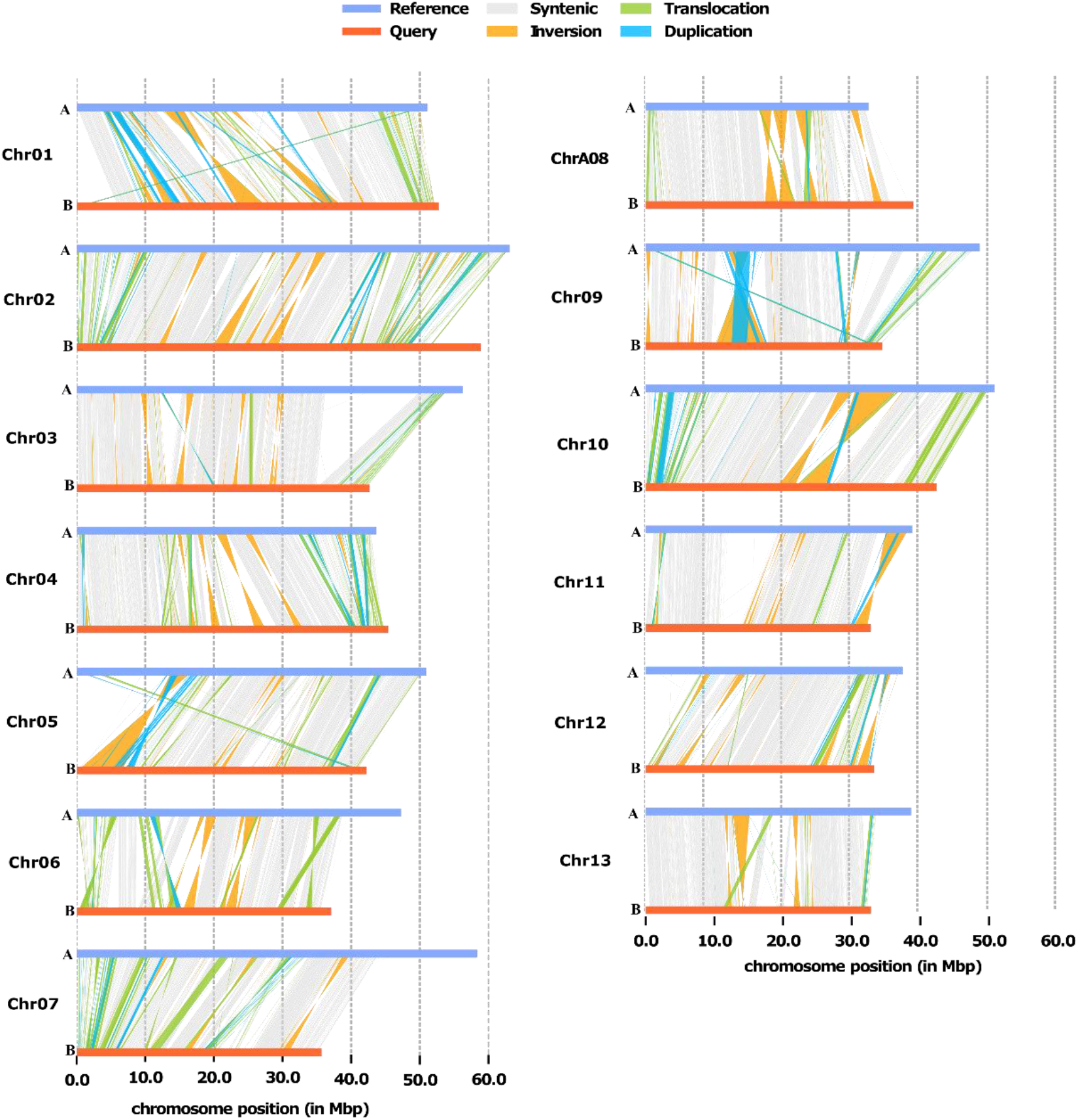
Structural variant analysis between A and B subgenomes. Chr01-13 shows the number of chromosomes in *T. sagittata* including the A subgenome (labeled with A) and B subgenome (labeled with B).

### Determination of secondary metabolite gene clusters and genes involved in the biosynthetic pathway BIA alkaloids in *T. sagittata*

PLANTISMASH v1.0 is a publicly available database to analyze secondary metabolite gene clusters (SMGCs) that are associated with biosynthesis of various compounds, such as alkaloids and terpenoids. We used this software to analyze potential SMGCs in the *T. sagittata* genome. This analysis obtained 58 putative SMGCs relating to different plant secondary metabolic pathways (Supplemental Data 7), which included seven alkaloids, three lignans, 20 saccharides, eight terpenes, five polyketides, and 15 genes putatively associated with biosynthesis of metabolites. The distribution of SMGCs were localized in all 26 chromosomes. Except for chromosomes A11, A12, B09, and B13, others have at least one SMGC. Data mining revealed that nine SMGCs consist of 82 genes, some of which encode methyltransferases.

To better understand whether these clusters were likely associated with biosynthetic pathways, we analyzed differential gene expression and co-expression networks with WGCNA to characterize DGEs. This analysis not only unearthed 17 clusters forming co-expression networks (Fig. S18-20, Data S8-9), but also clustered seven genes associated with the BIA biosynthetic pathway. In the assembled genomes, the seven genes are *TsA12G007000* (aspartate aminotransferase, mitochondrial), *TsB12G003350* (probable aminotransferase TAT2), *Tsatg001763lG000010* (*RS*-norcoclaurine 6-*O*-methyltransferase-like protein), *TsA13G009510* (3’-hydroxy-N-methyl-(*S*)-coclaurine 4’-*O*-methyltransferase 2-like), *TsA11G003380* (APapaver somniferum *O*-methyltransferase 1), and *TsA02G014550* ((*S*)-stylopine/(*S*)-canadine/(*S*)-nandinine synthase).

### Identification of key candidate genes involved in the jatrorrhizine biosynthesis

Enzymes including norcoclaurine synthase (NCS), different methyltransferases, and BIA bridge enzyme (BBE) have been proposed to catalyze steps from tyrosine to jatrorrhizine. BBE was proposed to convert (*S*)-reticuline to (*S*)-scoulerine through a methylene-bridge-forming reaction catalyzed by BBE. After *O*-methylation, a methylenedioxy bridge is added to the isoquinoline moiety by (*S*)-canadine synthase (CAS), a CYP719 protein (Li *et al*., 2020). Herein, our sequence mining and clustering analysis identified homologous gene copies encoding NCS (including both dioxygenase-like and pathogenesis-related 10-like protein genes), (*S*)-norcoclaurine 6-*O*-methyltransferase (6OMT), and (*S*)-canadine synthase (CAS) (Fig. 5A, Fig. S23, Data S11). More importantly, an analysis of homology searching and structural domain alignment (HMMER) identified 543 *CYP450* genes in the *T. sagittata* genome. The sequences of these genes were analyzed in the SwissProt database (Fig. S24, Data S12). Based on domain annotation and functional descriptions of homologs, the 543 genes were clustered into different families and subfamilies. In addition, a deep analysis of the sequences of members identify a CYP719 homolog, *TsA02G014550* (Data S12) localized on chromosome A02. Further phylogenetic analysis showed that *TsA02G014550* was clustered with CAS homologs from *P. somniferum* (XM_026596979.1), *C. japonica* CYP719A1 (AB026122.1), *and C. chinensis* (KC577598.1). These data suggest that *TsA02G014550* likely associates with the biosynthesis of jatrorrhizine.

**Fig. 5.**
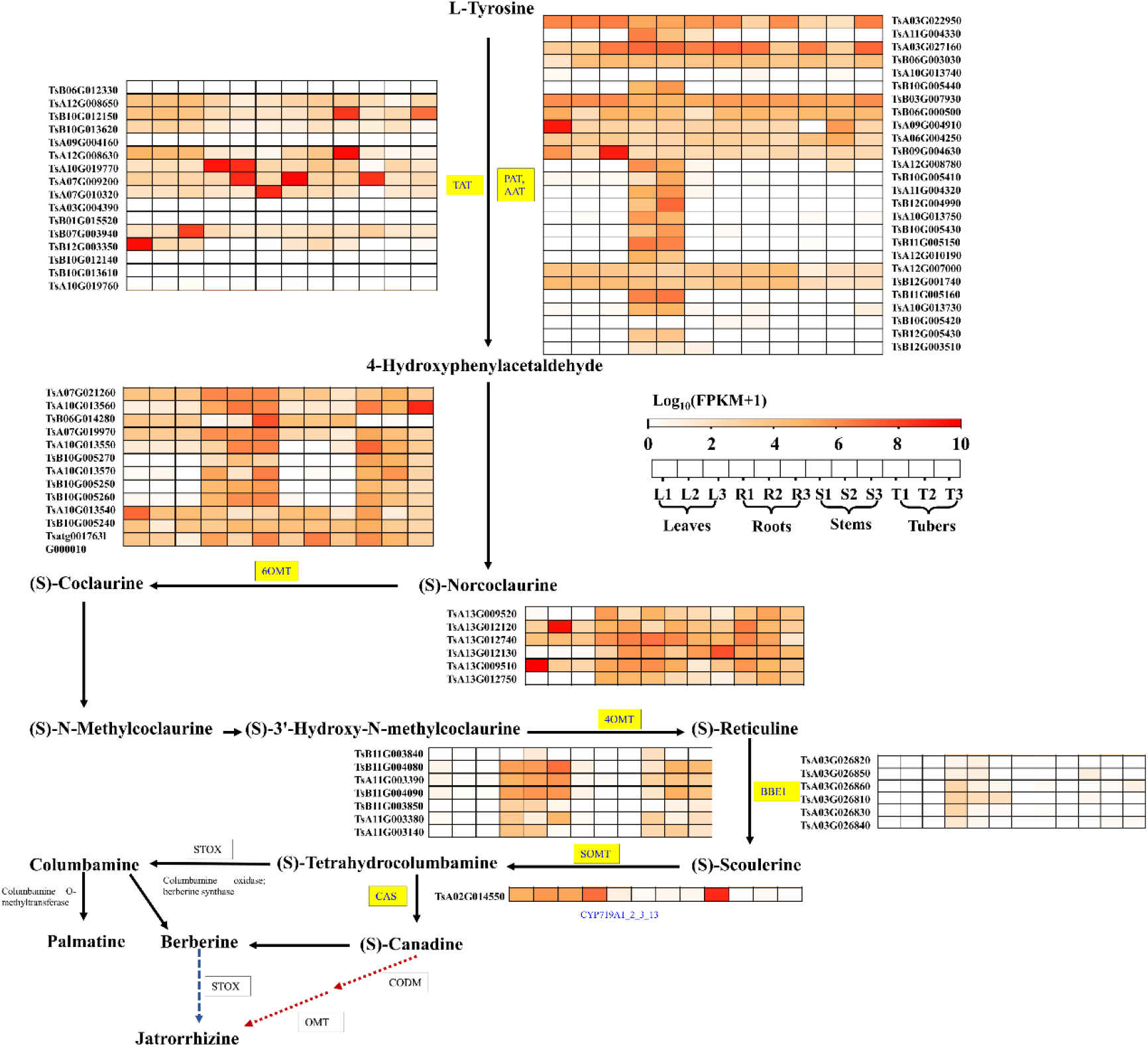
Mining of genes involved in the biosynthesis of BIA alkaloids in the *T. sagittata* genome. Enzymes associated with the biosynthetic pathway of jatrorrhizine are highlighted for each step. The expression profiles of gene encoding enzymes are indicated with heat maps.

In addition to targeted metabolic profiling described above, HPLC-MS was performed to further understand metabolite profiles in leaf, stem, root, and tuber tissues. Based on mass-to-charge ratio and fragmentation patterns, HPLC-MS analysis allowed annotating 124 alkaloids, including 32 isoquinoline alkaloids, 14 aporphine alkaloids, 23 phenolamine, 4 piperidine alkaloids, 10 plumerane, 3 pyridine alkaloids, and 37 other alkaloids (Fig. 6E, Fig. S22, and Data S13). Of the isoquinoline alkaloids, included were three enzymes, such as (S)-Norcoclaurine, methylcoclaurine, and scoulerine from the jatrorrhizine biosynthesis pathway. The abundance anlaysis determined 97 differentially accumulated alkaloids (DAAs) in four types of tissues. Four major DAA groups were 29 isoquinoline, 22 phenolamine, 11 aporphine, and 5 plumerane alkaloids (Data S14). Tissue pairs were compared to understand the specific accumulation of these DAAs. The resulting data showed, 64 from leaves versus tubers, 77 from stems versus tubers, and 48 from roots versus tubers. This comparison showed that the content of jatrorrhizine was the highest in leaves compared with stems, roots, and tubers (Fig. 6E).

**Fig. 6.**
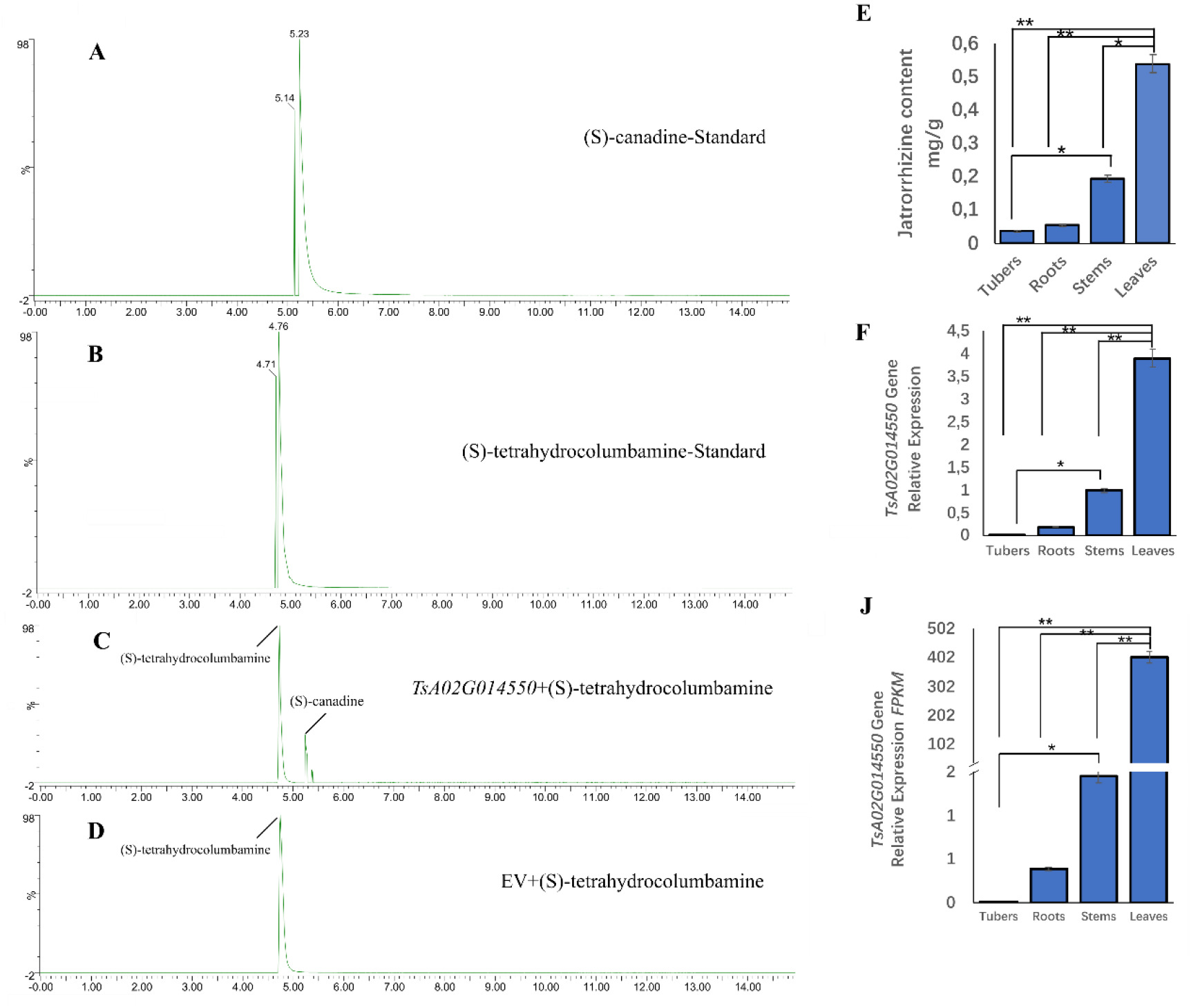
*In vitro* catalytic activity of a canadine synthase encoded *TsA02G014550* and its expression profiles. (A) (*S*)-canadine standard, (B) (*S*)-tetrahydrocolumbamine standard, (C) incubation n of recombinant protein induced from *TsA02G014550* gene and (*S*)-tetrahydrocolumbamine, (D) incubation of extraction of empty vector with (*S*)-tetrahydrocolumbamine, (E) The content of jatrorrhizine in different tissues of *T. sagittata* plants, (F) qRT-PCR data showed *TsA02G014550* gene expression profiles in different tissues, and (J) the *FPKM* values showed the *TsA02G014550* gene expression profiles in different tissues.

### In vitro catalytic analysis of *TsA02G014550*

Gene expression analysis via both qRT-PCT and RNA-seq showed that the expression level of *TsA02G014550* was higher in leaves than in other tissues (Fig. 6F and J). This result corresponded to the most abundance of jatrorrhizine in leaves. Based on this and the gene clustering analysis abovementioned, we proposed that *TsA02G014550* likely catalyzed a step in the biosynthetic pathway of jatrorrhizine. To test this hypothesis, the full-length cDNA of *TsA02G014550* was cloned into the pESC-His vector, which was transformed into the WAT11 yeast strain with modified endogenous NADPH-CYP reductase. A positive yeast colony was selected for *in vitro* assays using (*S*)-tetrahydrocolumbamine as a potential substrate (Fig. 6B). After this compound was feed to yeast, compounds were extracted for HPLC-MS analysis. This assay identified (*S*)-canadine produced from the feeding experiments, demonstrating that *TsA02G014550* encoded a (*S*)-canadine synthase (Fig. 6C).

## Discussion

*T. sagittata* is a polyploidy (allotetraploid) medicinal plant with a relatively large genome size (2.33 GB), which causes difficulties in genome sequencing. To overcome this challenge, we sequenced and assembled the genome of this plant with new sequencing technologies and assembly algorithms that were recently developed for the assembly of chromosome-scale and haplotype-resolved autopolyploid and allopolyploid plant genomes (Oh *et al*., 2020; Sun *et al*., 2022; Yang *et al*., 2019; Zhang *et al*., 2015). We integrated the PacBio long HiFi reads, Hi-C sequencing, and error corrections based on Illumina short reads data to assemble the *T. sagittata* genome at the chromosomal scale. Our genome assembly not only obtained the contig N50 of 8.3Mb and scaffold N50 of 42.06Mb, but also mapped 94.66% of the assembled sequences to 26 chromosomes (13 A sub-genome and 13 B sub-genome). Based on the mapping rate and quality of genome assembly (Korlach *et al*., 2017), our mapping rate indicated the continuity and completeness of the *T. sagittata* genome assembled. Furthermore, the analysis of BUSCO indicated 96.78% of gene completeness, supporting the high quality of the *T. sagittata* genome assembled here. In addition, our data showed 2.98% heterozygosity of the *T. sagittata* genome, supporting a common feature of the high heterozygosity discovered in other polyploidy plants (Miao *et al*., 2022; Shen *et al*., 2018; X. Xu *et al*., 2021; Yin *et al*., 2021).

It was interesting that approximately 51.72% of the *T. sagittata* genome consisted of repetitive elements. Our assembly observed that the *T. sagittata* genome was full of long terminal repeats (LTR) s distributed in the chromosomes in a nonrandom pattern. The characterization of repeats identified 239,772 and 211,805 intact *Gypsy* and *Copia* retrotransposons, respectively. This feature was also reported by the genomes of *C.chinensis* (Liu *et al*., 2021) and *T. wilfordii* (Tu *et al*., 2020), both of which were characterized to contain both *Gypsy* and *Copia* elements, with the former more than the later in abundance. Based on the observations from *C. chinensis* (Liu *et al*., 2021) and *T. wilfordii*, our data suggest that the two types of repetitive elements have played an important role in the genome evolution of *T. sagittata*, and the *Gypsy* repeats may have played more roles rather than the *Copia* elements. A phylogenomic analysis with *T. sagittata* and other 16 species revealed a close early-evolving eudicot placement of *T. sagittata* and *C. chinensis* (Fig. 2A). This datum supports that the roles of repetitive elements in the genome evolution of angiosperms.

Our assembly provides data to infer that an ancient WGD might have play an important role in the genome evolution of polyploidy. It has been documented that polyploidy events have occurred across land plants, contributed to their evolutionary success, and been playing a driving force in the acquirement of new functions of genes (Baurens *et al*., 2019). Like the genomes of opium poppy (*P. somniferum*) (Guo *et al*., 2018) and *C. chinensis* (Liu *et al*., 2021), we also observed an ancient WGD in the *T. sagittata* genome. To understand whether the ancient WGD played similar roles, a synteny dot plot was built with the *T. sagittata*, *P. somniferum* and *C. chinensis* genomes. This analysis of the plot revealed a 2:2 syntenic relationship among them and estimated that the ancient WGD in the *T. sagittata* genome occurred around 86 MYA. This syntenic analysis further suggested that the *T. sagittata* WGD events happened after the Menispermaceae-Papaveraceae-Ranunculaceae divergence. The similarity of the ancient WGDs between alkaloids-rich *T. sagittata* and *P. somniferum* suggests that such an ancient genome duplication has been likely associated with the diversity of BIA alkaloids in these and other plants. In particular, *PAT, TAT, CAS, BBE, 6OMT, 4OMT*, and *SOMT*, all were expanded to four copies in the *T. sagittata* genome. Gene duplication analysis indicated that the expansion of these pathway genes were duplicated from *P. somniferous* and *C. sinensis* in different periods (Fig. 3E and F). This gene duplication was supported by a family divergence analysis, which indicated the divergence of the Menispermaceae from Papaveraceae and Ranunculaceae about 107-120 MYA and 89-108 MYA, respectively. This result was agreement with a previous report that *P. somniferum* (Papaveraceae) was found to diverge from *Aquilegia coerulea* (Ranunculaceae) around 110 million years (Ma) ago (Guo *et al*., 2018). *CAS*, *BBE*, *6OMT*, *4OMT*, and *SOMT* encoding enzymes required for the biosynthesis of BIAs were proposed to be present in the common Ranunculales ancestor before the divergence of the Papaveraceae, Menispermaceae, and Ranunculaceae about 110 MYA (Li *et al*., 2020). These data propose that an ancient WGD leads to neofunctionalization and recruitment of additional enzyme classes responsible for the biosynthesis of BIAs.

The high quality of our assembly indicates that the *T. sagittata* genome enhances uncovering members of the CYP450 family associated with the biosynthesis of BIAs. It was reported that CYP-mediated methylenedioxy bridge formation (*CYP719* members) significantly contributed to the final diversity of protoberberine-type alkaloids (Mizutani and Sato, 2011). In our genome assembly, we mined the CYP450 family associated with the BIA and identified a candidate gene, *TsA02G014550* gene. Gene annotation indicated that this candidate was a *CAS* homolog. *In vitro* assay showed that its recombinant protein catalyzed (S)-tetrahydrocolumbamine to canadine (Fig. 6C). Similar observations were observed in other genome assemblies. *Cch00017825* from *C. chinensis* that was clustered with a *CAS* gene of *C. japonica CjCYP719A1* (99.11% identical) (Ikezawa *et al*., 2007) was found to encode methylenedioxy bridge-forming enzymes in the BIA biosynthesis pathway of *C. chinensis* (Liu *et al*., 2021). In enzyme reaction, the recombinant Cch00017825 only converted (*S*)-tetrahydrocolumbamine to (*S*)-canadine but also converted (*S*)-canadine to nandinine. A high quality genome was reported to be useful to isolate CYP450 genes from the genomes of California poppy (*E. californica*) and Mexican prickly poppy (*Argemone mexicana*). Three members, *CYP719A5*, *CYP719A2*, and *CYP719A3* were determined from the California poppy genome. *In vitro* assays showed that CYP719A5 has cheilanthifoline synthase activity (Ikezawa *et al*., 2009), while both CYP719A2 and CYP719A3 had a stylopine synthase activity (Ikezawa *et al*., 2007). More importantly, CYP719A3 was shown to use three substrates, (*R*, *S*)-cheilanthifoline, (*S*)-scoulerine, and (*S*)-tetrahydrocolumbamine. Similarly, *CYP719A13* identified in the Mexican prickly poppy (*Argemone mexicana*) was reported to encode a protein with a stylopine synthase activity (Díaz Chávez *et al*., 2011). These data suggest that gene duplication is associated with the evolution of new diverse functions.

In summary, a high quality genome of *T. sigattata* was assembled via short read (Illumina Hiseq) sequencing, long read sequencing (PabBio HiFi), and Hi-C sequencing. The genome is featured with a high heterozygosity and polyploidy. The assembled genome unearthed an ancient WGD event in *T. sagittata,* which was likely related to the divergence of Menispermaceae and Papaveraceae, and Ranunculaceae. The assembled genome allowed mining CYP450 family and obtained *TsA02G014550* that was a CAS homolog catalytically involved in the jatrorrhizine biosynthesis. Our findings provide new insights into the genome evolution in the Menispermaceae family and gene expansion associated with the biosynthesis of BIAs.

## Data availability

The data supporting findings of this work are available within the paper and its Supplementary Information files. A reporting summary for this article is available as a Supplementary Information file. The assembled genome have been deposited in NCBI under the BioProject accession number PRJNA942017.

## Acknowledgments

This work was supported by the Key Industries Innovation Chain Major Project, Hubei Province (2021ACA004, 2022AC003-01-003). We thank the Chinese Scholarship Council (CSC) for providing scholarships for our Ph.D. studies.

## Author contributions

A-MM, W-XK, Y-GZ, and S-SH conceived and designed the study. A-MM prepared the materials and conducted the experiments. A-MM, L-SB, O-Z, S-YH, and L-MJ analyzed the data and prepared the results. A-MM wrote the manuscript. S-SH, A-MM, Y-GZ, F-SQ, X-DY, M-ZN, and W-XK edited and improved the manuscript. All authors approved the final manuscript. A-MM and S-SH contributed equally to this work.

## Supporting information

### Figures

**Fig. S1.** Length distribution of PacBio long reads produced from the *T.sagittata* sample.

**Fig. S2**. Genome assembly flowchart demonstrating assembly polishing and data integration.

**Fig. S3.** The Hi-C heatmap of *T. sagittata* genome assembly.

**Fig. S4.** The gene annotation methods of *T. sagittata* genome.

**Fig. S5.** A species tree of 17 plant species based on the 1113 single copy orthologues using ASTRAL. All nodes with 100% support except denoted.

**Fig. S6.** Distribution of genes and gene families of plant species we investigated.

**Fig. S7.** Gene Ontology (GO) enrichment analysis of genes specifically presented in the *T. sagittata*.

**Fig. S8.** (KEGG) enrichment analysis of genes presented in the *T. sagittata*.

**Fig. S9.** Gene Ontology (GO) enrichment analysis of genes expanded in the *T. sagittata*.

**Fig. S10.** (KEGG) enrichment analysis of genes expanded in the *T. sagittata*.

**Fig. S11.** 4DTv analysis in *T. sagittata* genome.

**Fig. S12.** The relative age (Kimura distance) computed for LTR retroelements.

**Fig. S13.** Dot plot of synteny blocks between the *T. sagittata* genome and *Coptis chinensi*.

**Fig. S14.** Dot plot of synteny blocks between the *T. sagittata* genome and *Papaver somniferum*.

**Fig. S15.** SV detection length distribution

**Fig. S16.** SNP and InDel variation in reference A subgenome.

**Fig. S17.** PAV detection length distribution.

**Fig. S18.** Clustering dendrogram of differentially expressed genes with assigned, merged, and original module colors.

**Fig. S19.** Heatmap of the gene network using Topological Overlap Matrix among all differential expressed genes identified from different tissues.

**Fig. S20.** Heatmap of differentially expressed metabolites of *T. sagittata*.

**Fig. S21**. Transcription factors distribution in *T. sagittata* genome.

**Fig. S22.** Log_2_ relative content of alkaloids in *T. sagittata*.

**Fig. S23.** Phylogenetic trees of the genes involved in the Jatrorrhizine biosynthetic pathway.

**Fig. S24.** CYP450 genes distribution in different plant species.

### Tables

**Table S1.** Survey statistic results of *T. sagittata.*

**Table S2.** Sequencing data statistics of *T. sagittata*.

**Table S3.** Statistics of sequencing results.

**Table S4.** Reads length distribution statistics.

**Table S5.** Coverage statistics of *T. sagittata* genome.

**Table S6.** Statistics on the quantity of each type of Hi-C sequencing data.

**Table S7.** Hi-C assembly data statistics table.

**Table S8.** Assessment of the gene coverage rate using BUSCO.

**Table S9.** Assessment of the gene coverage rate using CEGMA.

**Table S10.** Primary statistical results of gene structure prediction of *T. sagittata* genome.

**Table S11.** Primary statistical results of gene structure prediction of *T. sagittata* and relative species.

**Table S12.** The statistical results of gene function annotation of *T. sagittata* genome.

**Table S13.** Non-coding RNA statistical results in *T. sagittata* genome.

**Table S14.** Pseudogene prediction results in *T. sagittata* genome.

**Table S15.** Motif annotation information statistics of *T. sagittata* genome.

**Table S16.** Repeat contents in the *T. sagittata* genome.

**Table S17.** Statistics of tandem repeats in *T. sagittata* genome.

**Table. S18.** Statistics on data mapping of RNA sequencing.

**Table S19.** Plant genomes for phylogenetic and comparative genomics analyses.

**Table S20.** Shows the number and percentage of genes involved in the collinearity analysis

**Table. S21.** The jatrorrhizine biosynthesis gene divergence from *C. chinensis.*

**Table. S22.** The jatrorrhizine biosynthesis gene divergence from *P. somniferum.*

**Table. S23.** Primer list of *TsA02G014550* gene.

**Table. S24.** Primer list of Actin gene.

### Data

**Data S1.** Summary of RNA sequencing.

**Data S2.** Gene family clustering statistical information table.

**Data S3.** Enriched Go terms for gene families specific to *T. sagittata.*

**Data S4.** Enriched KEGG terms for gene families specific to *T. sagittata.*

**Data S5.** Enriched GO terms for expanded genes in *T. sagittata*.

**Data S6.** Enriched KEGG terms for expanded genes in *T. sagittata*.

**Data S7.** SMGCs identified in *T. sagittata* genome.

**Data S8.** Genes classified into 17 modules in different colors by WGCNA.

**Data S9**. Differential expressed genes and their expression abundance matrix (fragments per kb exon model per million mapped reads) from 9 tissues of *T. sagittata*.

**Data S10.** TFs identified in *T. sagittata* genome.

**Data S11.** Berberine biosynthesis pathway genes expressed in different part *T. sagittata* plant.

**Data S12**. Enriched GO terms for CYPs genes in *T.sgaittata*.

**Data S13.** Type and relative content of alkaloids in different part of *T. sagittata*.

**Data S14.** Differentially accumulated alkaloids between different groups.

